# Trends and perspectives on the use of social network analysis in behavioural ecology: a bibliometric approach

**DOI:** 10.1101/379008

**Authors:** Quinn M.R. Webber, Eric Vander Wal

## Abstract

The increased popularity and improved accessibility of social network analysis has improved our ability to test hypotheses about the complexity of animal social structure. To gain a deeper understanding of the use and application of social network analysis, we systematically surveyed the literature and extracted information on publication trends from articles using social network analysis. We synthesize trends in social network research over time and highlight variation in the use of different aspects of social network analysis. Our primary finding highlights the increase in use of social network analysis over time and from this finding, we observed an increase in the number of review and methods of social network analysis. We also found that most studies included a relatively small number (median = 15, range = 4–1406) of individuals to generate social networks, while the number and type of social network metrics calculated in a given study varied zero to nine (median = 2, range 0–9). The type of data collection or the software programs used to analyze social network data have changed; SOCPROG and UCINET have been replaced by various R packages over time. Finally, we found strong taxonomic and conservation bias in the species studied using social network analysis. Most species studied using social networks are mammals (111/201, 55%) or birds (47/201, 23%) and the majority tend to be species of least concern (119/201, 59%). We highlight emerging trends in social network research that may be valuable for distinct groups of social network researchers: students new to social network analysis, experienced behavioural ecologists interested in using social network analysis, and advanced social network users interested in trends of social network research. In summary we address the temporal trends in social network publication practices, highlight potential bias in some of the ways we employ social network analysis, and provide recommendations for future research based on our findings.

## Introduction

Social network analysis is a powerful method used to measure social relationships among individuals (Wey, Blumstein, Shen, & Jordán, 2008) and it originates in mathematical graph theory, a body of literature on which many current practices are based (Biggs, Lloyd, & Wilson, 1976; Krause, Lusseau, & James, 2009). Social networks are composed of nodes (individuals) and edges (social connections between individuals), properties of a network can be used to quantify direct and indirect social relationships, which is the broad appeal of social network analysis. Despite the current perceived popularity and historical significance of social network analysis, there remains no systematic overview of social network analysis. We synthesize trends in social network analysis and highlight variation in publication trends over time with an aim to guide future research using social network analysis.

Social network analysis has been used since the 1950’s (for review see Krause, Lusseau, & James, 2009) and gained popularity among behavioural ecologists in two last decades. The emergence of network analysis to quantify social relationships has honed our questions and provided new avenues to test hypotheses about the causes and consequences of complex animal social structures (Croft, Madden, Franks, & James, 2011). As a result, animal social network analysis has become an important sub-discipline within behavioural ecology. Social behaviours, calculated using social network analysis, have been linked to a range of behavioural and ecological variables, including fitness (Stanton & Mann, 2012; Vander Wal, Festa-Bianchet, Réale, Coltman, & Pelletier, 2015), movement (Spiegel, Leu, Sih, & Bull, 2016), dominance (Bierbach et al., 2014), predation (Heathcote, Darden, Franks, Ramnarine, & Croft, 2017), animal personality (Wilson, Krause, Dingemanse, & Krause, 2013), information transfer (Firth, Sheldon, & Farine, 2016), pathogen dynamics (Webber et al., 2016), and quantitative genetics (Fisher & McAdam, 2017). Indeed, the application of social network analysis has become wide-spread.

Social network analysis involves three primary steps (*sensu* Farine & Whitehead, 2015). First, social association or interaction data are used to construct social networks. Animals can be observed interacting or associating (Altmann, 1974) or association can be inferred with bio-logging technology (for examples see Croft, Darden, & Wey, 2016). Second, social interaction or association data are converted into pairwise matrices and association indices are often calculated. This form of data conversion often involves correction of the data; for instance, heterogeneity in the number of observations per individual is corrected using the half-weight index (Cairns & Schwager, 1987). Third, statistical or mathematical modelling of social networks to test hypotheses about underlying social network structure. For instance, individual or group-level social network metrics may be generated and combined with attribute data (*sensu* Farine & Whitehead, 2015). Among the most important aspects of social network analysis is the ability to translate network-level social processes to individuals. A wide range of social network metrics exist (for definitions see Table 1; Silk et al., 2017; Wey et al., 2008), many of which are used as individual-based proxies for animal social interaction or association and can be used in statistical models. Despite the utility of network metrics for studies of individuals, most reviews on social network analysis do not explicitly provide guidance on the number or type of network metrics that should be quantified for a given type of analysis.

**Table 1:** Glossary of definitions for the most commonly used social network metrics and association indices. *x*: number of instances where individuals A and B were observed together; *Y_a_*: the number of instances where individual A was observed without vidual B; *Y_b_*: the number of instances where individual b was observed without individual A; *Y_ab_*: the number of instances where individuals A B that were observed at the same time but not together; *HWI_a_*: the half-weight index calculated for individual A; *HWI_b_*: the half-weight index ulated for individual B.

| Metric | Definition | Number of occurrences for each metric |
| --- | --- | --- |
| Degree | The number of direct connections an individual has with other individuals. Degree can be separated into in- and out-degree, where in-degree is the number of connections directed towards the focal individual and out-degree is the number of connections direct to non-focal individuals. | 97 (38 calculated in-degree; 36 calculated out-degree) |
| Graph strength | The combined weight (based on association indices or the frequency or duration of associations) of all of an individual's connections. | 97 (14 calculated in-strength; 12 calculated out-strength) |
| Clustering coefficient | The density of the subnetwork of a focal individual's neighbors, usually operationalized as the number of connections between neighbours divided by the possible maximum number of connections between them. | 87 |
| Betweenness centrality | The number of times an individual occurs on the shortest path-length between two other individuals in a network. | 68 (2 calculated in-betweenness) |
| Eigenvector centrality | The influence of an individual in a network based on the number and strength of the focal individuals connections. | 65 |
| Path length | The shortest number of edges that connect two individuals in a network. Path length is either the average, or sum, of all shortest number of edges between a focal individual and all other individuals in a network. | 24 |
| Closeness | The normalized mean path length from an individual to all other individuals. | 19 (4 calculated in-closeness; 4 calculated out-closeness) |
| **Group metrics** |  |  |
| Graph density | The proportion of realized edges in a network. | 58 |
| Modularity | Measure of the strength of division of a network into different modules, such as communities, clusters, and social groups. | 50 |
| Transitivity | The tendency for an individual's connections to associate with one another. | 13 |
| **Association Indices** |  |  |
| Half-weight index ($HWI_{ab}$) | $HWI_{ab} = \frac{x}{\frac{1}{2}(Y_a + Y_b) + Y_{ab} + x}$ | 66 |
| Simple ratio index ($SRI_{ab}$) | $SRI_{ab} = \frac{x}{x + Y_{ab} + Y_a + Y_b}$ | 65 |
| Gregarious-adjusted half-weight index ($HWIG_{ab}$) | $HWIG_{ab} = HWI_{ab} \frac{\sum HWI}{\sum HWI_a \cdot \sum HWI_b}$ | 5 |
| Twice-weight index ($AI_{A,B}$) | $AI_{A,B} = \frac{(Y_{ab})}{[(Y_A) + (Y_B) + (Y_{ab})]}$ | 4 |

Although social network analysis is an important method for testing hypotheses about animal social structure (Croft et al., 2011), it is also relevant in an applied context (Makagon, McCowan, & Mench, 2012; Snijders, Blumstein, Stanley, & Franks, 2017). Specifically, social network analysis has been used to quantify social structure for species of conservation concern as well as for captive and domestic species. In a killer whale (*Orcinus orca*) network, targeted removal of key individuals can fragment social networks and potentially reduce cohesiveness of highly dynamic social units (Williams & Lusseau, 2006). Moreover, social network analysis can also be used to predict pathogen dynamics (Drewe, 2010; Rushmore et al., 2013), which can have implications for reservoir hosts of infectious disease (Hamede, Bashford, Jones, & McCallum, 2012) or pathogen transmission from wild to domestic animals (Craft, 2015). Social network analysis of captive or domestic species also provides an opportunity to improve animal welfare and husbandry practices (Rose & Croft, 2015). Understanding social structure of captive and domestic species is important because many of captive species are highly gregarious and housed in social groups while in captivity. For example, understanding dominance-subordinate relationships between group members may be particularly important for captive species to reduce aggression and fighting (Makagon et al., 2012).

Despite the recent interest in social network analysis as a method, and subsequent emergence as a sub-discipline within behavioural ecology (Wey et al., 2008), there has been no systematic review of trends in social network publication. Here, we employ a bibliometric approach to synthesize research on animal social network analysis. Bibliometric approaches include objective measures of the content of a given research field. Recent examples within ecology and evolution include bibliometrics of conservation physiology (Lennox & Cooke, 2014), food web research (Tao et al., 2015), and disease ecology (Manlove et al., 2016). The intention of bibliometrics is to extract specific aspects of a research to assess, track, and analyze the status, gaps, and trends within a research field. As a method of evaluating research, bibliometrics are therefore intentionally non-prescriptive. By design, our bibliometric analysis is objective and is an empirical illustration of trends and gaps in social network research. Broadly, our four primary objectives were to:

1. Assess trends in publication on social network analyses over time and identify the types of journals where social network articles are published.
2. Describe social network methods from peer-reviewed articles, including the type of data collection methods and software used as well as the type of association indices and number of network metrics calculated.
3. Identify which species are being studied using social network analysis including conservation status and the potential for taxonomic bias.
4. 4) Based on some of our findings, we highlight four non-prescriptive directions for future research using social network analysis (Table 1).

## Methods

### Data collection

To evaluate the state of social network analysis within the peer-reviewed literature we conducted a literature survey using Thomson’s Scientific Web of Science. We used Web of Science because it allowed us to access large quantities of journal articles using thematic searches. We conducted our literature survey up to and including 2016 using a range of search terms that were likely to yield articles on social network analysis. We conducted four systematic searches of key-words on Web of Science which generated lists of articles that were likely relevant for our analysis. We searched the phrases ‘social network analysis’, ‘social network’, ‘network analysis’, and ‘contact network’. Searches were conducted between January 8^th^ and 21^st^, 2017. In total, we identified 1,603 unique articles through this method. We then used the ‘snowball approach’ to collect additional articles from the reference lists of all review and methodological articles (see below) identified through our Web of Science searches.

To meet the criteria for inclusion in our analysis we used a conservative and systematic filtering process. We only included articles that generated social networks based on pairwise social associations of non-human individual animals or, if empirical data were not included, discussed social network analysis in the form of a methodological, meta-analytical, or synthetic review (see below). We therefore excluded articles that generated networks where nodes represented, for example, spatial locations, or edges represented parasite sharing, while we also excluded ecological, genetic, and neural networks. Although these fields use similar techniques to social network analyses, their inclusion is beyond the scope of our analysis. We also excluded articles that modeled social network processes because these articles were not explicitly based on empirically-derived pairwise social associations, but, rather, the majority of articles that model social network dynamics are based on simulated data. In addition, we only included peer-reviewed research articles. We excluded peer-reviewed commentary and response articles, editorials, letters to the editor, prefaces to theme issues, conference proceedings, theses, book chapters, and books. Access to some of these sources of information can be sporadic and typical bibliometric analyses exclude these sources because it is difficult to systematically search grey literature. From our original output of 1,603 articles only 293 articles matched our criteria. The snowball approach yielded an additional 134 articles for a total of 427 relevant peer-reviewed articles that used or where about social network analysis that met our criteria. While it is possible we missed some published articles about social network analysis, we are confident that we identified a very large proportion of the literature up to 2016 in an unbiased manner, which accurately represents the trends in social network analysis implementation. All data collected are available as supplementary material.

### Data extraction and analyses

To address our first objective, we assessed temporal changes in the number of articles published using, or about, social network analyses per year. We first assigned articles to one of four categories: empirical, review, methodological, or meta-analysis. Empirical articles contained data collected in the field or laboratory that were used to generate social networks; review articles were data-free and outlined a broad synthetic or theoretical contribution to social network analysis; methodological articles may, or may not, have contained empirical data, but provided an overview or demonstration of a particular methodological aspect of social network analysis; meta-analyses were categorized as articles that contained summaries or comparisons of empirically collected social network data from multiple species. We also assigned all articles to one of six general themes based on the journal of publication: behavioural (e.g. *Animal Behaviour*), general biology (e.g. *Proceedings of the Royal Society B*), general ecology (e.g. *Journal of Animal Ecology*), taxa-specific (e.g. *Journal of Fish Biology*), applied (e.g. *Applied Animal Behaviour Science*), or disease/parasitology journals (e.g. *Journal of Wildlife Diseases*). We recorded the year of publication for each article to demonstrate possible temporal changes in publication trends (see below).

To address our second objective, we extracted methodological aspects of social network articles. Our methodological overview comprised of four key aspects of social network analyses. First, we determined the software used to generate social network analysis. Second, we assessed how networks were constructed and how many networks were constructed in each article. We determined whether an association index was used, and, if not, whether network edges were weighted using another measure, such as frequency or duration of social interactions. We also counted the number of networks generated in each article as well as the number of uniquely identifiable individuals within each network. Third, to consider the technological aspect of social network analysis, we recorded how data were collected to generate networks and the general type of behaviour used to construct social networks. Finally, we summarized the statistical aspect of social network analyses by recording the number and type of social network metrics quantified in each article as well as the number of individuals in each network. We separated social network metrics into individual and group-level metrics. We used chi-square tests to test for potential differences in the proportion of articles using a given software, how data were collected, behaviours that were measured, association indices, and the type of network metrics.

To address our third objective, we extracted information on the study species and the broad taxonomic group of the study species from empirical articles. For each species we obtained IUCN listing as well as the taxonomic class of each species in our dataset.

## Results

### Objective 1

We identified 427 social network articles from the peer-reviewed literature and classified 337 (79%) as empirical, 52 (12%) as synthetic reviews, 27 (6%) as methodological, and 11 (3%) as meta-analytical (Figure 1). The first article in our database was from 1999 and we observed an exponential increase in the total number of social network articles over time (Figure 1).

**Figure 1.**
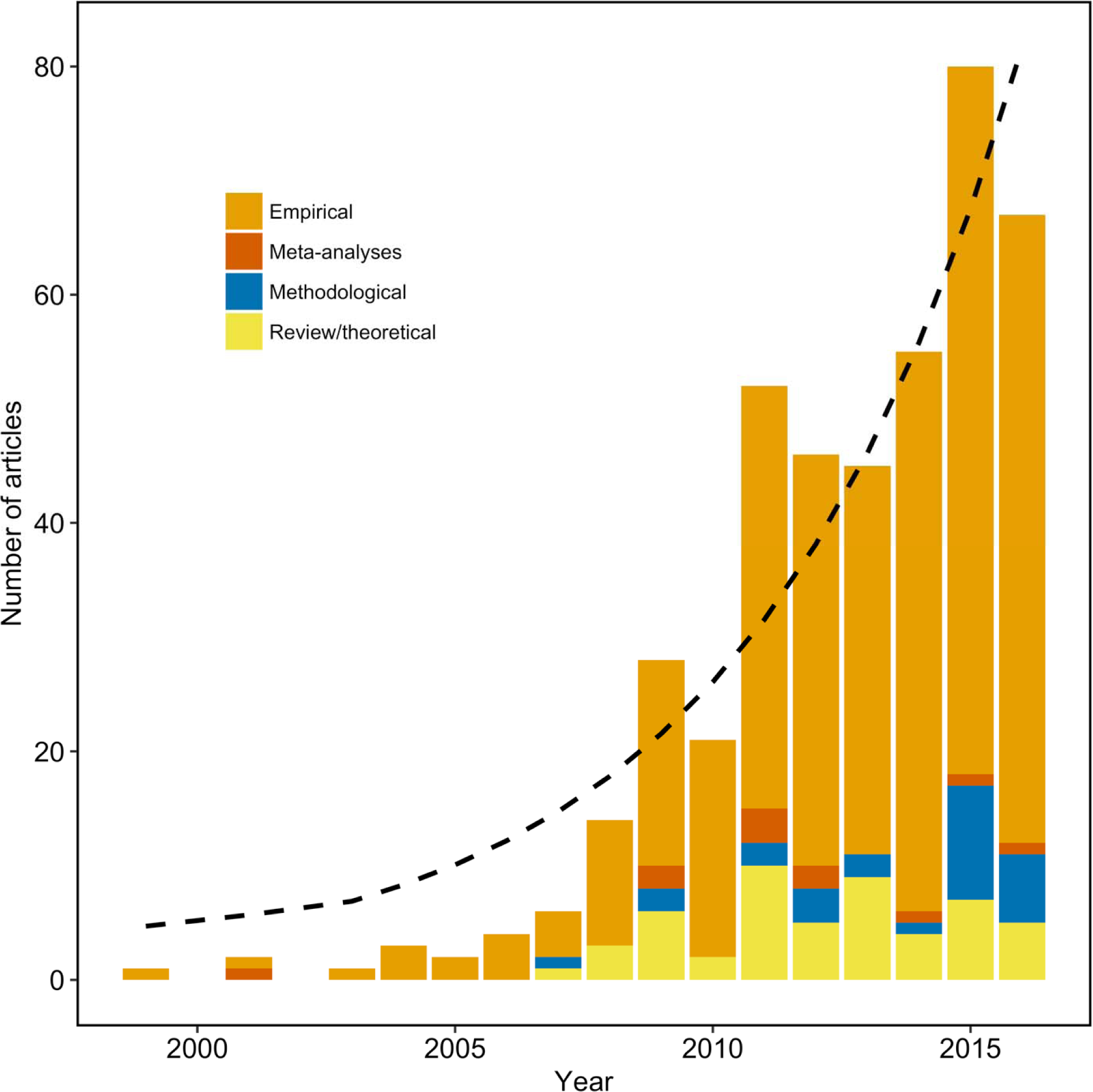
Number of social network articles over time separated by article type. Note, the dashed line represents an exponential growth curve for the number of social network articles published each year.

Social network articles were not published equally among journal types published (χ^2^ = 212.5, df = 5, p < 0.001). We found significant difference in the types of journal in which social network articles are. The most common type of journal where social network studies were published was behavioural journals (163/427, 38.2%), while social network studies were also published in general biology (125/427, 29.3%), taxa-specific (67/427, 15.7%), general ecology (44/427, 10.3%), applied (22/427, 5.1%), and disease/parasitology (6/427, 1.4%) journals (Figure 2).

**Figure 2.**
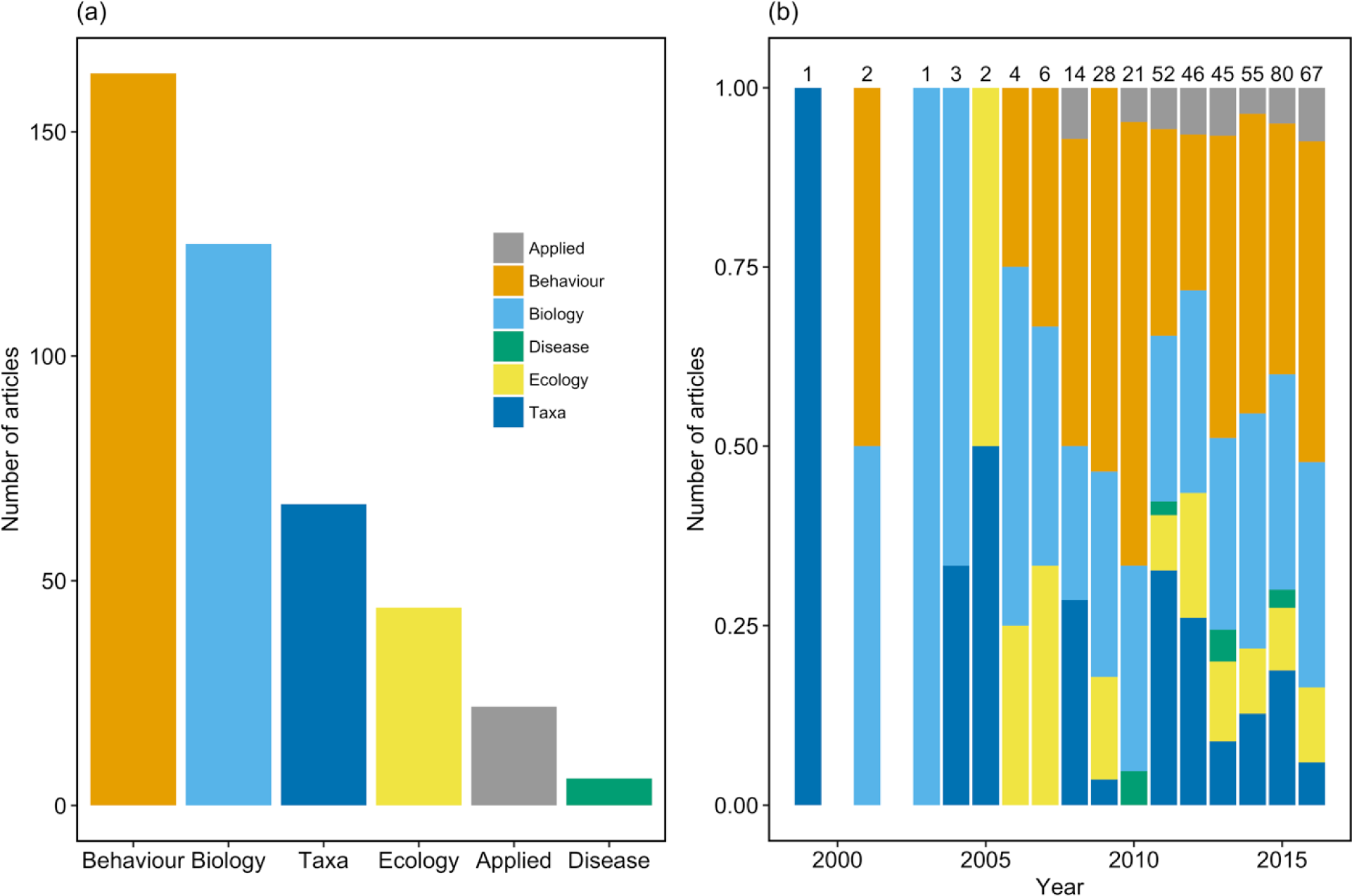
(a) Number of social network articles published in journals allocated to one of six categories. Note, social network articles are most commonly published in behavioural or general biology journals, while they are rarely published in journals categorized as applied or disease/parasitism. (b) Proportion of social network articles published in journals allocated to one of six categories over time. Note, number of articles for each year are provided.

### Objective 2

For empirical articles, the number of networks calculated per article was right-skewed, with a median of 3 networks per articles (SD = 15.1, range: 1–128, Figure 3a). The number of individuals in a given network was also right-skewed, with a median of 15 individuals per network (SD = 101, range = 4–1406, Figure 3b). The median number of social network metrics calculated per article was 2.0 (SD = 2.0, range: 0–9, Figure 3c), while the median number of network metrics per article that calculated at least 1 metric was 3.0 (SD = 1.8. range: 1–9).

**Figure 3.**
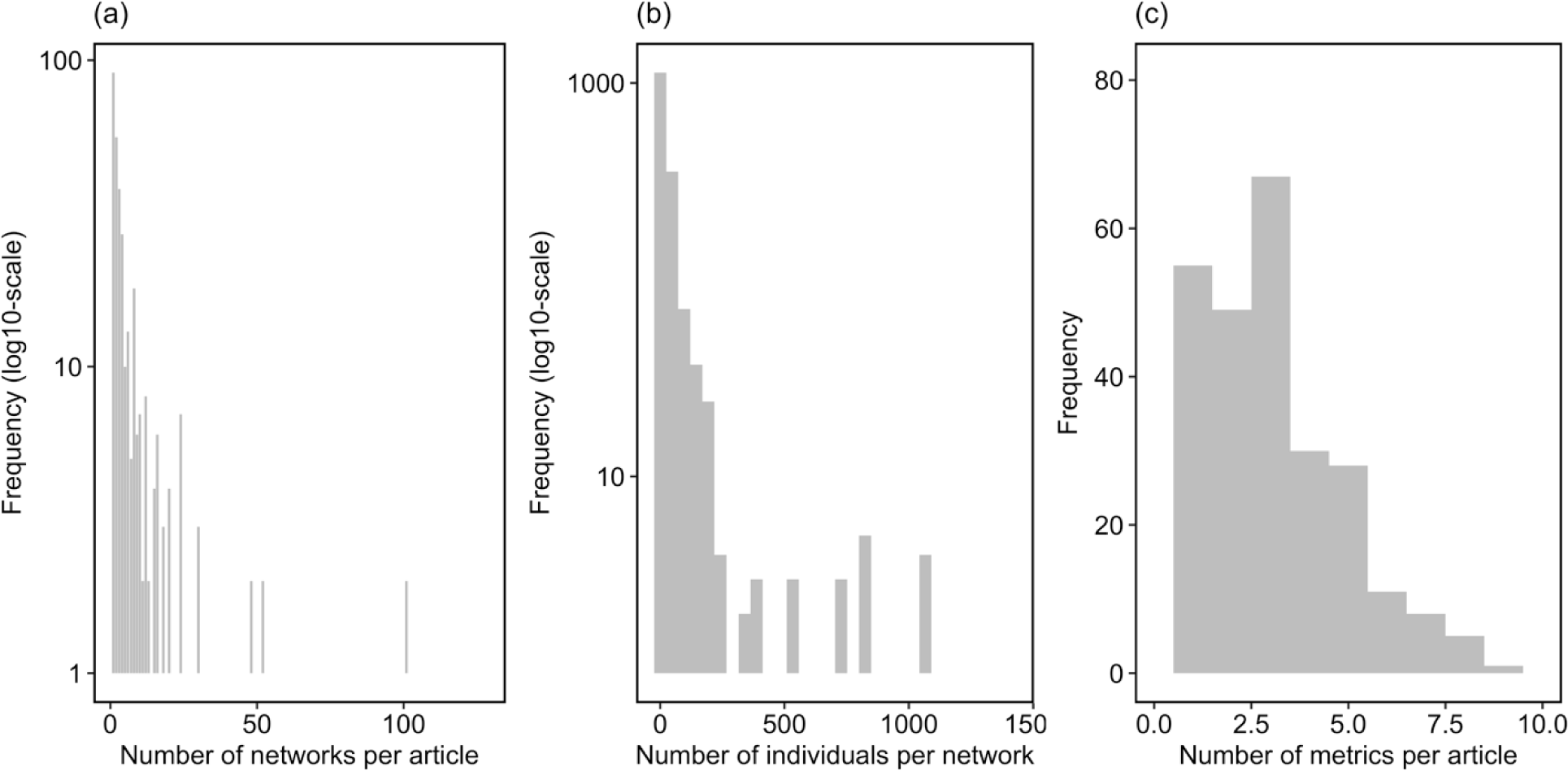
(a) Number of social networks generated per empirical article included in our study; median number of articles generated per article was 3 (range: 1–128). (b) Number of individuals per network generated; median number of individuals per network was 15 (range: 4–1406) from a total of 2674 networks generated across all 337 empirical articles. (c) Number of social network metrics generated per empirical article; the median number of metrics per article was 2 (range: 0–9).

In total, at least one social network metric was calculated in 254 of the 337 empirical articles we identified. Of the 254 studies that calculated at least one metric, we identified 35 unique social network metrics, and we found that network metrics were not used equally in articles where metrics were calculated (χ^2^ = 1297.1, df = 34, p < 0.001). The most commonly quantified individual-level social network metrics were degree and graph strength (both quantified in 97/254 articles, 38.2%), while clustering coefficient (87/254, 34.3%), betweenness centrality (68/254, 26.7%), and eigenvector centrality (65/254, 25.6%) were also commonly quantified. Meanwhile, the most commonly quantified group-level social network metrics were graph density (58/254, 22.8%) and modularity (50/254, 19.7%).

The type of software program used to generate social network was not equal across articles (χ^2^ = 33.3, df = 5, p < 0.001). The most commonly used software program was R (75/337 of empirical studies, 22.4%), including ‘*asnipe’*, ‘*igraph*’, and ‘*sna*’ among others (Figure 4), while SOCPROG (63/337 empirical studies, 18.8%) and UCINET (51/337 empirical studies, 15.2%) were the second and third most commonly used software programs, while 18.2% (61/337) articles used multiple software programs and 5% (20/337) articles used other software programs (Figure 4). We were unable to determine what software program was used in 64/337 (19%) of empirical articles.

**Figure 4.**
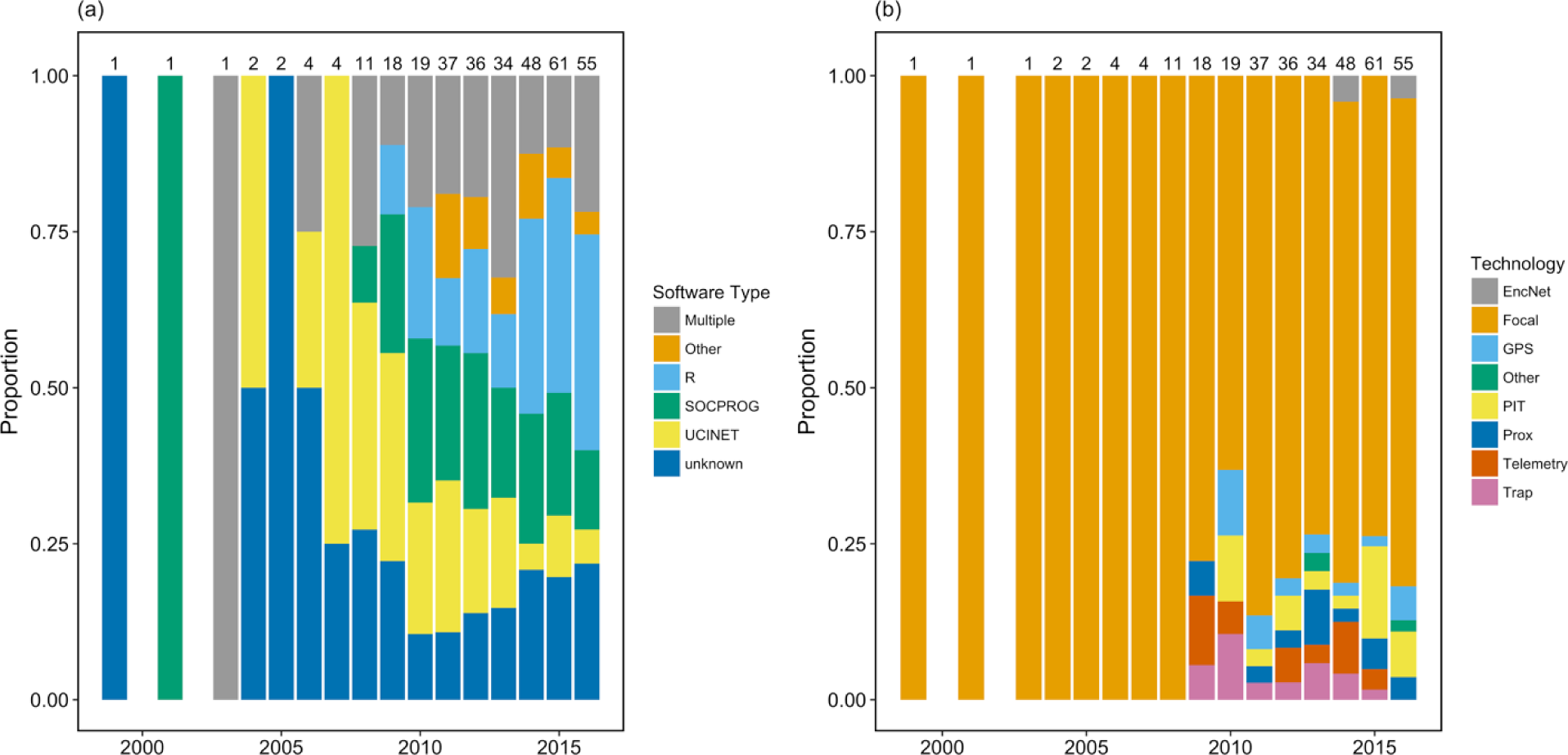
(a) Proportion of empirical social network articles that used different software programs to generate social networks over time. Note, most articles use at least one R package, although many articles use one of SOCPROG, UCINET, or some combination of R, SOCPROG, and UCINET. (b) Proportion of empirical social network articles which use different data collection techniques over time (EncNet: Encounter Net; GPS: global positioning system; PIT: passive integrated transponders; Prox: proximity devices; Trap: trapping). Note, the most common data collection technique is focal observation of animals, while a range of different technologies were used to varying degrees. Total numbers of empirical articles are presented at the top of each bar.

Data collection techniques differed among social network articles (χ^2^ = 1345, df = 7, p < 0.001). The most common data collection technique was focal observation (263/337, 78%, Figure 3b). A range of bio-logging technologies, including passive integrated transponders (20/337. 5.9%), proximity devices (12/337, 3.6%), VHF telemetry (12/337, 3.6%), GPS telemetry (11/337, 3.3%), trapping data (10/337, 3.0%), encounter-net (4/337, 1.1%) and other techniques (2/337, 0.5%), were also used to collect data (Figure 3b).

The type of association indices used to generate social networks differed among articles (χ^2^ = 254.7, df = 6, p < 0.001). The most common type of matrix weighting was based on the duration or frequency of interactions (132/337, 39.4%), while the HWI was the most commonly used association index (66/337, 19.7%), followed by the SRI (62/337, 18.5) the HWI_G_ (5/337, 1.4%) and the TWI (3/337, 0.1%), while in 45/337 (14.3%) articles, the association index or matrix weighting procedure was unknown, and 20/337 (5.9%) of articles used a binary association matrix.

The type of data collection methods used to describe associations were similarly employed among social network articles (χ^2^ = 1.3, df = 2, p = 0.52). Gambit-of-the-group was the most commonly used type of association data (119/337, 35.3%), while proximity to conspecifics (113/337, 33/3%) and behavioural interaction (102/337, 30.2%) were also relatively common.

#### Objective 3

In total, 201 unique species from 12 taxonomic classes were studied in empirical social network articles. Species were not equally studied within the IUCN red list categories (χ^2^ = 484, df = 8, p < 0.001). The most commonly observed listing was least concern (119/201, 59%), while species that were not listed (22/201, 11%), vulnerable (15/201, 7.5%), endangered (14/201, 7.0%), not threatened (12/201, 6.0%), critically endangered (7/201, 3.5%), data deficient (5/201, 2.5%), and domestic (4/201, 2.0%) were less commonly studied (Figure 5).

**Figure 5.**
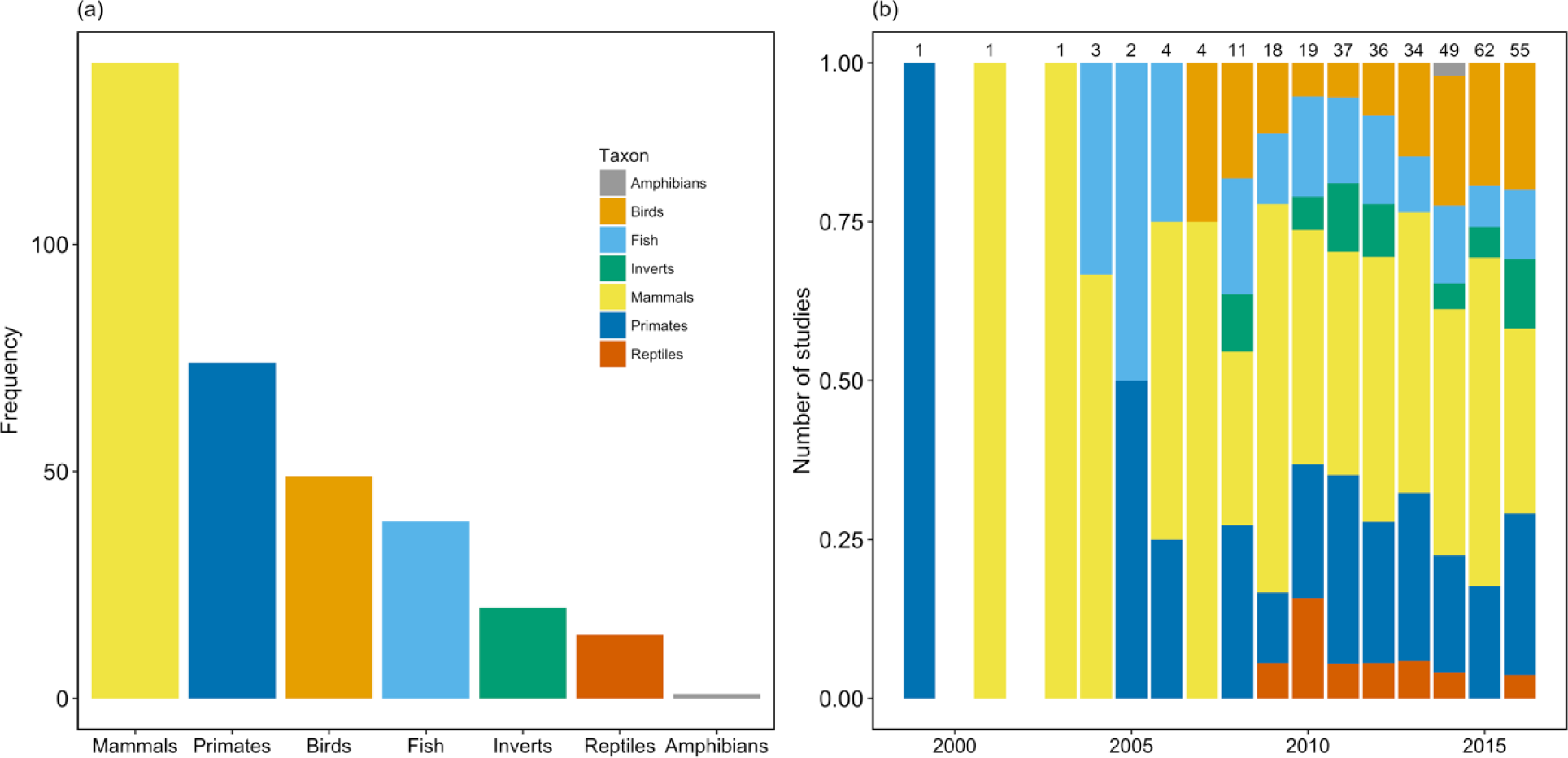
(a) Number of species from empirical social network articles from one of seven broad taxonomic groups, where the colours representing each taxonomic group in this panel carry over to panel (b). Primates were separated from non-primate mammals because of the large number of social network studies on primates. Note, most species studied in social network articles are either least concern or not listed, and the majority of these species are non-primate mammals or birds. (b) Proportion of species from empirical social network articles separated by broad taxonomic group over time. Note, the trends in species are similar over time with few observable discrepancies in the proportion of a given taxonomic group studied at any given time.

The proportion of species from different taxonomic classes was not equal among social network articles (χ^2^ = 690, df = 11, p < 0.001). The majority of species studied were in classes mammalia (111/201, 55%) and aves (47/201, 23%), while fewer were from classes actinopterygii (14/201, 7%), insecta (12/201, 6%), chondrichthyes (5/201, 2.5%), and reptilia (4/201, 2%). Meanwhile, there was a single (1/201, 0.005%) species studied in classes hymenoptera, malacostraca, sauropsida, amphibia, and arachnida. The most common species studied in all empirical social network articles were *Parus major* (17/337 empirical articles, 5%), *Poecilia reticulata* (14/337, 4.2%), *Pan troglodytes* (13/337, 3.9%), *Macaca mulatta* (12/337, 3.6%), *Tursiops truncatus* (11/337, 3.3%), and *Tiliqua rugosa* (10/337, 3.0%).

## Discussion

Social network analysis has increased in use over the last 20 years, and within this growing body of literature we found a lack of consensus regarding the number, and type, of network metrics calculated across studies, the use of association indices, and the species studied. We found that most empirical social networks contain relatively few individuals, while the use of specific association indices varies considerably across studies. We also found taxonomic bias in the species being studied, where the majority of species studied were birds or mammals and of these, most were listed least concern by the IUCN. After two decades of social network analysis, our study provides insight into some trends of network research within the field of animal behavioural ecology.

### Objective 1: trends in publication analysis

It is clear the use of social network analysis has increased in popularity over the past two decades, a trend which is broadly consistent with the general increase in academic publication. The number of empirical studies increased slowly between 1998–2007 before an exponential increase in the number of articles per year after 2008. Similar to many natural populations, there could be a carrying capacity for the number of articles using social network analysis in a given year, however, we may still be far from the actual carrying capacity of social network articles. Interestingly, the number of review and methods articles did not follow the same proportional increase as empirical articles, although in the final two years of our analysis (2015 and 2016) there was a qualitative increase in the number of methodological articles published. The recent increase in methods articles may be a function of a field that was largely missing a coherent and up to date set of guidelines. Prior to 2015, when a number of important methods articles were published (e.g., Farine & Whitehead, 2015; Whitehead & James, 2015), few comprehensive ‘how to’ articles explicitly provided guidance on social network analysis (although comprehensive books did exist: Croft et al., 2008; Whitehead, 2008). Importantly, an increase in methodological articles, as well as reviews about certain methods highlights an attempt to achieve a higher standard for social network methods up to the end of our data collection in 2016 (see references above) as well as in the short time since (e.g., Hoppitt & Farine, 2018; Silk & Fisher, 2017).

### Objective 2: methods

The advancement, availability, and accessibility of computer programming has undoubtedly contributed to the rise in popularity of social network analysis over time. Until the advent of open access software and relatively easy-to-use social network R packages (e.g., Csárdi & Nepusz, 2006; Farine, 2013, 2014) our findings suggest the social network sub-discipline appeared to have gone through some degree of methodological disarray. The increase in the proportion of methods articles published in 2015 and 2016 may have stimulated the formation, and implementation, of guidelines for social network analysis. For example, the selection and implementation of association indices is an important example of social network methodology that appears to lack general consensus. Accounting for the frequency or duration social associations, as well as the possibility of missing individuals during observation (Hoppitt & Farine, 2018; Silk, 2017), by using an association index should be standard operating procedure. By contrast, our analysis suggests that many studies do not use association indices. One possible explanation is the vast majority of empirical social network studies are generated using data collected by focal observation. Focal observations of individual behaviour may be more conducive for observing all individuals in a group, a scenario which does not necessarily require an association index if all individuals are monitored at the same time. Despite this possible explanation, it remains important for authors to ensure they follow appropriate methodological procedures. We recommend Farine and Whitehead (2015) as the most valuable and constructive outline of social network methods. Although our results suggest little consistency in the use of association indices in social network research, the recent publication of several important reviews on the topic suggest potential for future consensus.

The use of different social network software has also changed over time. SOCPROG and UCINET were widely used in the first decade of social network analysis. More recently, R packages, including *asnipe*, *igraph*, and *sna* have been more commonly used, while more specialized packages are currently in development (*ant*: Sosa et al., 2018). The implementation of social network analysis in R makes it more compatible with other analytical techniques, such as quantitative genetics (Thomson, Winney, Salles, & Pujol, 2018; Wilson et al., 2010), geographic patterns of space use (Spiegel et al., 2016), exponential random graph models (Silk & Fisher, 2017), or multi-layered social networks (Silk, Finn, Porter, & Pinter-Wollman, 2018). As social network analysis becomes more widely integrated within ecology we expect the use of R packages to generate social networks will continue to increase while the use of SOCPROG, UCINET, and other Windows or MatLab based programs are decreasing over time, potentially to a point where they are no longer used.

We also found the vast majority of empirical social network studies collected data by focal observation. Although focal observation is the most accurate method of collecting social association or interaction data, the use of biologging devices can increase the volume social network data. The most popular biologging device used to generate social network data is radio-frequency identification devices (RFIDs), such as passive integrated transponder tags, although most articles using RFIDs are from the same system (e.g., Aplin et al., 2013). We expect the availability of various types of biologging devices should increase the popularity of remotely collected social network data. For example, GPS telemetry data and autonomous fixed arrays are relatively under-used technologies in social network research, despite the fact that both have potential to test a variety of hypotheses about animal social structure (Jacoby & Freeman, 2016). Proximity devices represent a promising data collection method (Prange, Jordan, Hunter, & Gehrt, 2006), although in the ~10 years since their development they also remain relatively under-used in social network research. Potential explanations include issues with data processing and the reliability of detection distance among devices (for discussion see Boyland, James, Mlynski, Madden, & Croft, 2013; O’Brien, Webber, & Vander Wal, 2018). We expect the use of various biologging technologies, including GPS telemetry, autonomous fixed arrays, proximity devices, and RFIDs, to conduct social networks will continue to increase in usage over time.

Social network metrics are arguably the most appealing aspect of network analysis because of they can be interpreted in a traditional context and used in statistical models. We observed apparent preference for certain social network metrics. Graph strength and degree were the most commonly calculated social network metrics, while clustering coefficient, betweenness and eigenvector centrality were common. Among the review and methods articles that we identified there is little discussion or guidance on the number or type network metrics to calculate in a given study. While we do not advocate for the use of any particular network metrics, the use of a large number of social network metrics in a given study may be problematic. A potential implication of calculating multiple network metrics without *a priori* justification may violate assumptions of most frequentist models (see Table 2). A potential solution is the *a priori* selection of biologically relevant network metrics and justification for using a given network metric or the use of multiple competing hypotheses to compare models using different network metrics (Table 2).

**Table 2.** Outline of implications, opportunities, and future directions for future research based on key findings from our study.

| **Finding** | **Implication** | **Opportunities and future directions** | **Key references** |
| --- | --- | --- | --- |
| **Number of individuals per network (Figure 3b)** | Potential for bias if a small proportion of the group or population is included as well as less variation in apparent social phenotypes in small networks. | At minimum, studies should justify the use of smaller proportions of individuals, but when possible, we recommend the inclusion of as many individuals as possible in empirical networks. | Hoppitt & Farine, (2018) |
| **Number of network metrics per article (Figure 3c)** | Depending on the type of analysis conducted, measuring multiple network metrics with *a priori* justification may violate assumptions of most frequentist models. | The use of multiple competing hypotheses can reduce the violations of frequentist statistics and improve clarity of studies calculating network metrics. | — |
| **Software program used for network analysis (Figure 4a)** | Increase in the use of various R packages and decrease in less flexible programs. | The use of R to generate social networks represents a seamless and reproducible way to integrate output from social network analysis with other forms of statistical or mathematical analyses. | Farine, (2013); Jacoby & Freeman, (2016) |
| **Data collection method (Figure 4b)** | While focal observation is a reliable form of data collection, the use of bio-logging technology increases ability to measure social connections at certain times (e.g. at night or in winter) and for certain species (e.g. small or cryptic species). | Low prevalence of most bio-logging technologies in social network research represents an opportunity to test hypotheses about the ecology and evolution of social structure using existing datasets. | Prange et al., (2006); Boyland et al., (2013) |

### Objective 3: Species

Conservation is becoming an increasingly important issue across ecological disciplines. Among behavioural ecologists, understanding conservation implications for species is often cited as an important conclusion of empirical work. Meanwhile, several recent reviews have also highlighted the relative complacency of behavioural ecologists in a conservation context (Caro & Sherman, 2011, 2013) and our findings generally support these views. The field of ‘conservation behaviour’ (Blumstein, 2010; Macdonald, 2016) has highlighted social network analysis as a tool to help inform management plans, particularly for highly gregarious species in the wild (Snijders, Blumstein, Stanley, & Franks, 2017). Our findings suggest that most existing studies using social network analysis are conducted on mammals and birds of least concern.

Little is known about the relationship between social behaviour and population dynamics and social network analysis represents a potential method that could improve our understanding of this relationship (Webber & Vander Wal, 2018). Allee effects, a phenomenon described as a positive relationship between fitness and group size or population density (Stephens & Sutherland, 1999), are an important conservation issue for some social species (Angulo et al., 2017). Social networks could provide information about group or population level phenomenon that could inform our understanding of Allee effects. In addition, for highly gregarious species, individuals with high centrality may have higher fitness (Stanton & Mann, 2012; Vander Wal et al., 2015) and for species of conservation concern, the use of social network analysis could improve management of groups or populations. Moreover, social network analysis may be a useful tool to monitor groups of translocated individuals to ensure individuals maintain social structure or are able to integrate with new group members (e.g., Poirier & Festa-Bianchet, 2018). Indeed, social network analysis has potential to inform conservation practices.

The most common species in our dataset were model species from well-established systems, most of which are relatively abundant and easy to work with. The use of model species in established systems almost certainly improves our understanding of the ecology and evolution of social behaviour, and in support of this line of research, the majority of species in our database were listed by the IUCN as either least concern, data deficient, domestic, or not listed. By contrast, some studies highlight conservation implications for their results, even though the species being studied are not listed, and the relevance of social network analysis only applies for a relatively small proportion of species in our dataset. As is the case for methods being applied to managed populations, social network analysis is likely most beneficial when applied directly to a given population, while there may be limited applicability across species or even populations.

## Conclusion

We systematically and objectively synthesized trends in social network research using a bibliometric approach. Social network analysis has increased exponentially over time, while there is a wide range of data collection and analytical methods used in studies that generate social networks. Our assessment of social network methods suggests there is little consensus in the number, and type, of network metrics calculated across studies, the use of association indices, and the species studied. Despite the potential impact of social network analysis for species of conservation concern, a relatively small proportion of empirical studies examined species listed above least concern. In summary, we highlight emerging trends in social network research that may be valuable for three distinct groups of social network researchers. First, students new to social network analysis will benefit from our synthesis of the usage of different social network methods. Second, experienced behavioural ecologists interested in using social network analysis will benefit from our synthesis because we highlight technical aspects of network analysis, including an approximate minimum number of individuals to be included in a network as well as different data collection strategies used in network research, details which may be relevant for potential social network users to design data collection strategies. Third, advanced social network users will benefit from the knowledge of the wide variation in usage for some trends in social network research and the identification of certain practices or aspects of network research that are becoming more, or less, common. Social network analysis has come a long way in the last two decades and as theoretical and practical aspects of network analysis further develop, we anticipate increased consensus on usage, methods, and applications within behavioural ecology.

## Acknowledgements

We thank C. Prokopenko, J. Aubin, P. O’Brien, A. Robitaille, and S. Zabihi-Seissan for excellent comments on that improved the manuscript. Funding for was provided by a Vanier Canada Graduate Scholarship to QMRW and a National Sciences and Engineering Research Council Discovery Grant to EVW.

## References

Altmann, J. (1974). Observational study of behavior: Sampling methods. Behaviour, 49(3), 227–267.

Angulo, E., Luque, G. M., Gregory, S. D., Wenzel, J. W., Bessa-Gomes, C., Berec, L., & Courchamp, F. (2017). Allee effects in social species. Journal of Animal Ecology, (October 2016), 1–12. http://doi.org/10.1111/1365-2656.12759

Aplin, L. M., Farine, D. R., Morand-Ferron, J., Cole, E. F., Cockburn, A., & Sheldon, B. C. (2013). Individual personalities predict social behaviour in wild networks of great tits (Parus major). Ecology Letters, 16(11), 1365–1372. http://doi.org/10.1111/ele.12181

Bierbach, D., Oster, S., Jourdan, J., Arias-Rodriguez, L., Krause, J., Wilson, A. D. M., & Plath, M. (2014). Social network analysis resolves temporal dynamics of male dominance relationships. Behavioral Ecology and Sociobiology, 68(6), 935–945. http://doi.org/10.1007/s00265-014-1706-y

Biggs, N., Lloyd, E. K., & Wilson, R. J. (1976). Graph Theory, 1736-1936. Oxford University Press.

Blumstein, D. T. (2010). Social behaviour in conservation. Social Behaviour, 520–534. http://doi.org/10.1017/CBO9780511781360.041

Boyland, N. K., James, R., Mlynski, D. T., Madden, J. R., & Croft, D. P. (2013). Spatial proximity loggers for recording animal social networks: Consequences of inter-logger variation in performance. Behavioral Ecology and Sociobiology, 67(11), 1877–1890. http://doi.org/10.1007/s00265-013-1622-6

Cairns, S. J., & Schwager, S. J. (1987). A comparison of association indices. Animal Behaviour, 35, 1454–1469.

Caro, T., & Sherman, P. W. (2011). Endangered species and a threatened discipline: Behavioural ecology. Trends in Ecology and Evolution, 26(3), 111–118. http://doi.org/10.1016/j.tree.2010.12.008

Caro, T., & Sherman, P. W. (2013). Eighteen reasons animal behaviourists avoid involvement in conservation. Animal Behaviour, 85(2), 305–312. http://doi.org/10.1016/j.anbehav.2012.11.007

Craft, M. E. (2015). Infectious disease transmission and contact networks in wildlife and livestock. Philosophical Transactions of the Royal Society of London. Series B, Biological Xciences 370(1669), 1–12. http://doi.org/10.1098/rstb.2014.0107

Croft, D. P., Darden, S. K., & Wey, T. W. (2016). Current directions in animal social networks. Current Opinion in Behavioral Sciences 12, 52–58. http://doi.org/10.1016/j.cobeha.2016.09.001

Croft, D. P., Madden, J. R., Franks, D. W., & James, R. (2011). Hypothesis testing in animal social networks. Trends in Ecology and Evolution, 26(10), 502–507. http://doi.org/10.1016/j.tree.2011.05.012

Croft, D. P., Ruxton, G. D., & Krause, J. (2008). Exploring Animal Social Networks. Princeton University Press.

Csárdi, G., & Nepusz, T. (2006). The igraph software package for complex network research. InterJournal Complex Systems 1695, 1–9.

Drewe, J. A. (2010). Who infects whom? Social networks and tuberculosis transmission in wild meerkats. Proceedings of the Royal Society B 277(1681), 633–42. http://doi.org/10.1098/rspb.2009.1775

Farine, D. R. (2013). Animal social network inference and permutations for ecologists in R using asnipe. Methods in Ecology and Evolution, 4(12), 1187–1194. http://doi.org/10.1111/2041-210X.12121

Farine, D. R. (2014). Measuring phenotypic assortment in animal social networks: Weighted associations are more robust than binary edges. Animal Behaviour 89, 141–153. http://doi.org/10.1016/j.anbehav.2014.01.001

Farine, D. R., & Whitehead, H. (2015). Constructing, conducting and interpreting animal social network analysis. Journal of Animal Ecology 84, 1144–1163. http://doi.org/10.1111/1365-2656.12418

Firth, J.., Sheldon, B. ., & Farine, D. R. (2016). Pathways of information transmission among wild songbirds follow experimentally imposed changes in social foraging structure. Biology Letters, 12(6), 20160144. http://doi.org/10.1098/rsbl.2016.0144

Fisher, D. N., & McAdam, A. G. (2017). Social traits, social networks and evolutionary biology. Journal of Evolutionary Biology, 1–16. http://doi.org/10.1111/jeb.13195

Hamede, R., Bashford, J., Jones, M., & McCallum, H. (2012). Simulating devil facial tumour disease outbreaks across empirically derived contact networks. Journal of Applied Ecology, 49(2), 447–456. http://doi.org/10.1111/j.1365-2664.2011.02103.x

Heathcote, R. J. P., Darden, S. K., Franks, D. W., Ramnarine, I. W., & Croft, D. P. (2017). Fear of predation drives stable and differentiated social relationships in guppies. Scientific Reports 7(February), 41679. http://doi.org/10.1038/srep41679

Hoppitt, W. J. E., & Farine, D. R. (2018). Association indices for quantifying social relationships: how to deal with missing observations of individuals or groups. Animal Behaviour 136, 227–238. http://doi.org/10.1016/j.anbehav.2017.08.029

Jacoby, D. M. P., Croft, D. P., & Sims, D. W. (2012). Social behaviour in sharks and rays: Analysis, patterns and implications for conservation. Fish and Fisheries, 13(4), 399–417. http://doi.org/10.1111/j.1467-2979.2011.00436.x

Jacoby, D. M. P., & Freeman, R. (2016). Emerging Network-Based Tools in Movement Ecology. Trends in Ecology & Evolution, xx. http://doi.org/10.1016/j.tree.2016.01.011

Krause, J., Lusseau, D., & James, R. (2009). Animal social networks: An introduction. Behavioral Ecology and Sociobiology, 63(7), 967–973. http://doi.org/10.1007/s00265-009-0747-0

Lennox, R., & Cooke, S. J. (2014). State of the interface between conservation and physiology: a bibliometric analysis. Conservation Physiology, 2, 1–9. http://doi.org/10.1093/conphys/cou003.Introduction

Macdonald, D. W. (2016). Animal behaviour and its role in carnivore conservation: examples of seven deadly threats. Animal Behaviour 120, 197–209. http://doi.org/10.1016/j.anbehav.2016.06.013

Makagon, M. M., McCowan, B., & Mench, J. A. (2012). How can social network analysis contribute to social behavior research in applied ethology? Applied Animal Behaviour Science, 138(3–4, SI), 152–161. http://doi.org/10.1016/j.applanim.2012.02.003

Manlove, K. R., Walker, J. G., Craft, M. E., Huyvaert, K. P., Joseph, M. B., Miller, R. S., … Cross, P. C. (2016). “One Health” or Three? Publication Silos Among the One Health Disciplines. PLOS BIOLOGY, 14(4). http://doi.org/10.1371/journal.pbio.1002448

O’Brien, P. P., Webber, Q. M. R., & Vander Wal, E. (2018). Consistent individual differences and population plasticity in network-derived sociality: An experimental manipulation of density in a gregarious ungulate. PLoS ONE, 13(3), e0193425.

Poirier, M.-A., & Festa-Bianchet, M. (2018). Social integration and acclimation of translocated bighorn sheep (Ovis canadensis). Biological Conservation 218(November 2017), 1–9. http://doi.org/10.1016/j.biocon.2017.11.031

Prange, S., Jordan, T., Hunter, C., & Gehrt, S. D. (2006). New radiocollars for the detection of proximity among individuals. Wildlife Society Bulletin, 34(5), 1333–1344.

Rose, P., & Croft, D. (2015). The potential of Social Network Analysis as a tool for the management of zoo animals. Animal Welfare, 24(2), 123–138. http://doi.org/10.7120/09627286.24.2.123

Rushmore, J., Caillaud, D., Matamba, L., Stumpf, R. M., Borgatti, S. P., & Altizer, S. (2013). Social network analysis of wild chimpanzees provides insights for predicting infectious disease risk. Journal of Animal Ecology, 82(5), 976–986. http://doi.org/10.1111/1365-2656.12088

Silk, M. J. (2017). The next steps in the study of missing individuals in networks: a comment on Smith et al. (2017). Social Networks. http://doi.org/10.1016/j.socnet.2017.05.002

Silk, M. J., Croft, D. P., Delahay, R. J., Hodgson, D. J., Boots, M., Weber, N., & McDonald, R. A. (2017). Using social network measures in wildlife disease ecology, epidemiology, and management. BioScience, 67(3), 245–257. http://doi.org/10.1093/biosci/biw175

Silk, M. J., Finn, K. R., Porter, M. A., & Pinter-Wollman, N. (2018). Can Multilayer Networks Advance Animal Behavior Research? Trends in Ecology & Evolution, 1–3. http://doi.org/10.1016/j.tree.2018.03.008

Silk, M. J., & Fisher, D. N. (2017). Understanding animal social structure: exponential random graph models in animal behaviour research. Animal Behaviour 132, 137–146. http://doi.org/10.1016/j.anbehav.2017.08.005

Snijders, L., Blumstein, D. T., Stanley, C. R., & Franks, D. W. (2017). Animal social network theory can help wildlife conservation. Trends in Ecology & Evolution, 32(8), 567–577. http://doi.org/10.1016/j.tree.2017.05.005

Sosa, S., Puga-Gonzalez, I., Hu, F. E., Zhang, P., Xiaohua, X., & Sueur, C. (2018). A multilevel statistical toolkit to study animal social networks: Animal Network Toolkit (ANT) R package. BioarXiv, (June). http://doi.org/10.1101/347005

Spiegel, O., Leu, S. T., Sih, A., & Bull, C. M. (2016). Socially interacting or indifferent neighbours? Randomization of movement paths to tease apart social preference and spatial constraints. Methods in Ecology and Evolution 7, 971–979. http://doi.org/10.1111/2041- 210X.12553

Stanton, M. A., & Mann, J. (2012). Early Social Networks Predict Survival in Wild Bottlenose Dolphins. PLoS ONE, 7(10), 1–6. http://doi.org/10.1371/journal.pone.0047508

Stephens, P. A., & Sutherland, W. J. (1999). Consequences of the Allee effect for behaviour, ecology and conservation. Trends in Ecology & Evolution 14, 400–401.

Tao, J., Che, R., He, D., Yan, Y., Sui, X., & Chen, Y. (2015). Trends and potential causations in food web research from a bibliometric analysis. Scientometrics, 105(1), 451–463. http://doi.org/10.1007/s11192-015-1679-2

Thomson, C. E., Winney, I. S., Salles, O. C., & Pujol, B. (2018). A guide to using a multiple-matrix animal model to disentangle genetic and nongenetic causes of phenotypic variance. BioRxiv. http://doi.org/http://dx.doi.org/10.1101/318451

Vander Wal, E., Festa-Bianchet, M., Réale, D., Coltman, D. W., & Pelletier, F. (2015). Sex-based differences in the adaptive value of social behavior contrasted against morphology and environment. Ecology, 96(3), 631–641. http://doi.org/10.1890/14-1320.1.sm

Webber, Q. M. R., Brigham, R. M., Park, A. D., Gillam, E. H., O’Shea, T. J., & Willis, C. K. R. (2016). Social network characteristics and predicted pathogen transmission in summer colonies of female big brown bats (Eptesicus fuscus). Behavioral Ecology and Sociobiology, 70(5), 701–712. http://doi.org/10.1007/s00265-016-2093-3

Webber, Q. M. R., & Vander Wal, E. (2018). An evolutionary framework outlining the integration of individual social and spatial ecology. Journal of Animal Ecology 87, 113–127. http://doi.org/10.1111/1365-2656.12773

Wey, T., Blumstein, D. T., Shen, W., & Jordán, F. (2008). Social network analysis of animal behaviour: a promising tool for the study of sociality. Animal Behaviour, 75(2), 333–344. http://doi.org/10.1016/j.anbehav.2007.06.020

Whitehead, H. (2008). Analyzing animal societies: quantitative methods for vertebrate social analysis. University of Chicago Press.

Whitehead, H., & James, R. (2015). Generalized affiliation indices extract affiliations from social network data. Methods in Ecology and Evolution, 6, 836–844. http://doi.org/10.1111/2041-210X.12383

Williams, R., & Lusseau, D. (2006). A killer whale social network is vulnerable to targeted removals. Biology Letters, 2(May), 497–500. http://doi.org/10.1098/rsbl.2006.0510

Wilson, A. D. M., Krause, S., Dingemanse, N. J., & Krause, J. (2013). Network position: A key component in the characterization of social personality types. Behavioral Ecology and Sociobiology, 67(1), 163–173. http://doi.org/10.1007/s00265-012-1428-y

Wilson, A. J., Réale, D., Clements, M. N., Morrissey, M. M., Postma, E., Walling, C. A., … Nussey, D. H. (2010). An ecologist’s guide to the animal model. Journal of Animal Ecology, 79(1), 13–26. http://doi.org/10.1111/j.1365-2656.2009.01639.x

